# High resolution cryo-EM of V-ATPase in native synaptic vesicles

**DOI:** 10.1101/2024.04.01.587493

**Authors:** Claire E. Coupland, Ryan Karimi, Stephanie A. Bueler, Yingke Liang, Gautier M. Courbon, Justin M. Di Trani, Cassandra J. Wong, Rayan Saghian, Ji-Young Youn, Lu-Yang Wang, John L. Rubinstein

**Affiliations:** Molecular Medicine Program, The Hospital for Sick Children, Toronto, Canada, M5G 1X8; Department of Medical Biophysics, The University of Toronto, Toronto, Canada, M5G 1L7; Department of Biochemistry, The University of Toronto, Toronto, Canada, M5S 1A8; Lunenfeld-Tanenbaum Research Institute, Toronto, Canada, M5G 1X5; Neuroscience and Mental Health Program, The Hospital for Sick Children, Toronto, Canada, M5G 1X8; Department of Physiology, The University of Toronto, Toronto, Canada, M5S 1A8; Cell Biology Program, The Hospital for Sick Children, Toronto, Canada, M5G 1X8; Department of Molecular Genetics, The University of Toronto, Toronto, Canada, M5S 1A8

## Abstract

Neurotransmitter release into the synaptic cleft between neurons enables the intercellular communication central to the function of the brain. In the presynaptic neuron, the proton pumping vesicular- or vacuolar-type ATPase (V-ATPase) powers neurotransmitter loading into synaptic vesicles (SVs), with the V_1_ complex dissociating from the membrane region of the enzyme before exocytosis. We isolated SVs from rat brain using SidK, a V-ATPase-binding bacterial effector protein. Single particle electron cryomicroscopy of the vesicles allowed high-resolution structure determination of V-ATPase within the native SV membrane. In the structure, regularly spaced cholesterol molecules decorate the enzyme’s rotor and the abundant SV protein synaptophysin binds the complex stoichiometrically. Conditions where ATP hydrolysis drives glutamate loading result in loss of V_1_ from the SV membrane, suggesting that SV loading is sufficient to induce V-ATPase dissociation.

## Introduction

The computational power of the brain arises from the ability of neurons to communicate with each other. This communication occurs across the chemical synapse formed between an axon terminal from one neuron and a dendrite from another. Arrival of an action potential at the axon terminal allows calcium to flow into the cytosol, stimulating neurotransmitter release into the synaptic cleft and activating postsynaptic receptors on the dendrite^1^. Within the axon terminal, neurotransmitters are stored within ∼40 nm diameter organelles known as synaptic vesicles (SVs), which go through a cycle of loading, exocytosis, and endocytosis for recycling (**Fig. 1A**). SV loading with neurotransmitter is powered by the vacuolar- or vesicular-type ATPase (V-ATPase), which establishes a transmembrane electrochemical proton gradient that is used by transport proteins to translocate neurotransmitters from the cytosol to the SV lumen. Following loading but before exocytosis, the V_1_ region of V-ATPase dissociates from the rest of the complex^2,3^. The SV membrane contains numerous proteins, including the vesicular-SNARE (SNAp REceptor) synaptobrevin-2 and the calcium-sensor synaptotagmin, which are essential for engaging the target-SNAREs syntaxin and SNAP-25 on the presynaptic membrane for SV exocytosis.

**Figure 1.**
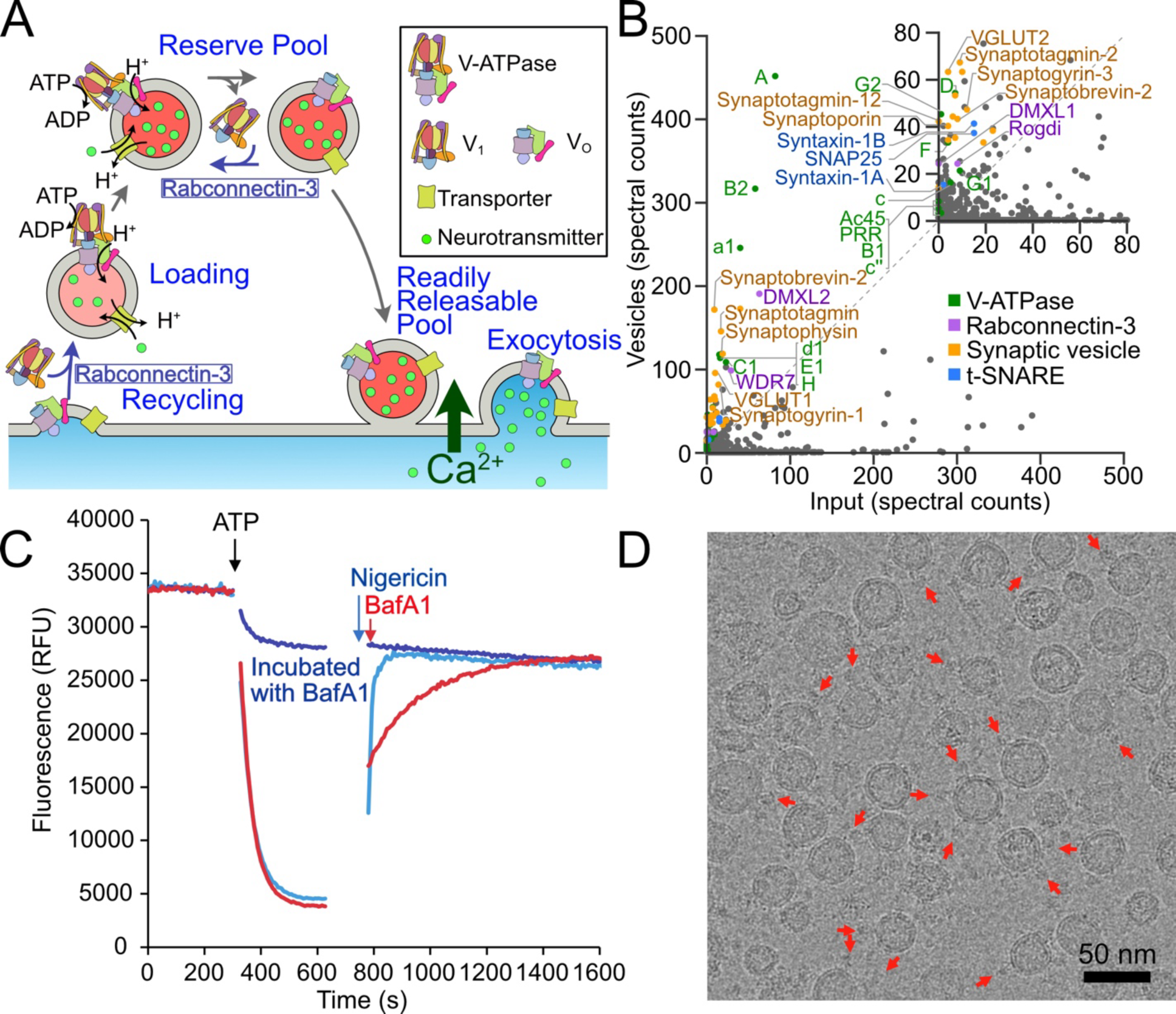
Isolation of SVs. **A,** SV cycle in the axon terminal. **B,** Mass spectrometry analysis of SVs and the rat brain lysate from which they were isolated. The scatter plot shows summed spectral counts from two preparations of each sample, one processed with trypsin and the other with chymotrypsin. **C,** SVs demonstrate acidification in an ACMA fluorescence-quenching assay. RFU, relative fluorescence units. **D,** Cryo-EM image of SVs. Red arrows indicate example V-ATPase complexes.

The mammalian V-ATPase is an assembly of sixteen different proteins^4^. There are multiple isoforms for many of these subunits, encoded by different genes^5^. The SV enzyme comprises subunits A, B2, C1, D, E1, F, G2, H, a1, c, c″, d1, e2, f (RNAseK), ATP6AP1/Ac45, and ATP6AP1/PRR^6^. ATP hydrolysis in the A_3_B_3_ subcomplex of the V_1_ region of the enzyme drives rotation the rotor (subunits DFc_9_c″, ATP6AP1/Ac45, and ATP6AP2/PRR). This rotation leads to proton translocation across the membrane through the interface of subunit a1 and the c-ring within the membrane embedded V_O_ region of the complex. Three peripheral stalk structures, each consisting of a pair of EG subunits, holds V_1_ stationary relative to V_O_ during proton pumping. V-ATPases are regulated by reversible dissociation of V_1_ from V_O_^7,8^, with ATP hydrolysis inhibited in the isolated V_1_ complex^9^. The structure of detergent-solubilized V-ATPase from mammalian brain was determined by electron cryomicroscopy (cryo-EM)^6,10^. Although V-ATPase forms numerous protein-protein interactions in other acidified organelles^11^ it is not known if or how the enzyme interacts with the rest of the SV machinery as it has not been observed in its native context. To understand the structure and function of endogenous V-ATPase within native SV membranes, we isolated ∼40 nm diameter vesicles from rat brain using the affinity of the *Legionella pneumophila* effector protein SidK for the V-ATPase. Mass spectrometry showed that the vesicles are SVs, and functional assays demonstrate that they are capable of ATP-driven acidification. Three-dimensional (3D) reconstruction of V-ATPase directly from images of the SVs revealed that V-ATPase forms a stoichiometric complex with the SV protein synaptophysin, and that treating the vesicles with a buffer that causes loading induces some of the V_1_ complexes to dissociate from the membrane.

## Results

### SidK enables rapid isolation of vesicles containing V-ATPase from rat brain

Multiple purification procedures have been described for the isolation of SVs from mammalian brain^12,13^. However, we developed a new procedure based on the affinity of the *Legionella pneumophila* effector protein SidK to V-ATPase^14^. Recombinant SidK modified with a 3×FLAG affinity tag previously allowed isolation of detergent-solubilized V-ATPase from rat brain^6^ and subsequently from cultured human cells^15^, porcine kidney^16^, and even from lemons^17^. SidK binding inhibits V-ATPase activity by ∼30%^18,19^, but despite this inhibition the rapid purification of V-ATPase afforded by SidK provides an enzyme preparation with specific activity approximately twice that of enzyme isolated by conventional chromatography^10,16^. Therefore, we homogenized a rat brain, removed cell debris by centrifugation, and passed the homogenate through an anti-FLAG affinity column charged with a 3×FLAG-tagged fragment of SidK. Negative stain electron microscopy showed that the material eluted from the column consisted of vesicles and free V_1_ complexes (**Fig. S1A**). These vesicles were subsequently separated from the V_1_ complexes by gel filtration chromatography (**Fig. S1B**). As described previously^20^, we found that homogenizing the whole brain to immediately free SVs, rather than first forming synaptosomes, led to the best yield of vesicles.

### Mass spectrometry shows that the purified vesicles are SVs

To characterize the protein content of the purified vesicles, we performed mass spectrometry of the vesicles and the centrifuged rat brain lysate from which the vesicles were isolated (i.e. the input material for the purification). Samples were digested separately with trypsin and with chymotrypsin, the latter helping to identify hydrophobic proteins that often lack trypsin cleavage sites. Comparison of total spectral counts for the fully purified samples versus the input material reveals enrichment of numerous SV markers (**Fig. 1B**, *orange*, **Supplementary Data 1**). These markers include the neurotransmitter transporters VGLUT1 and 2; the vesicular inhibitory amino acid transporter Slc32a1; several isoforms of the v-SNARE synaptobrevin; synaptotagmin; the abundant SV protein synaptophysin; and the synaptic vesicle glycoproteins 2A, 2B, and 2C. No lysosomal markers are enriched in the preparation, indicating that the vesicles are predominantly SVs and not derived from other cellular compartments where V-ATPase is abundant. The sample also contains the V-ATPase subunit isoforms expected in SVs (**Fig. 1B**, *green*): subunits A, B2, C1, D, E1, F, G2, H, a1, c, c″, d1, ATP6AP1/Ac45, and ATP6AP2/PRR. Only the small and hydrophobic V-ATPase subunits e2 and f were not detected. Subunit G1 was also detected, consistent with the earlier finding that subunit G1 makes up ∼20% of the G subunits in detergent-solubilized V-ATPase purified from rat SVs^6^. Components of Rabconnectin-3, which is responsible for reassembly of V_1_ with V_O_ following dissociation, were also detected in the vesicle preparation (**Fig. 1B**, *purple*). Rabconnectin-3 consists of subunits Rabconnectin-3α (DMXL2/1), Rabconnectin-3β (WDR7), and the proposed component Rogdi^21^, all of which are enriched in the sample. The yeast equivalent of Rabconnectin-3, RAVE (<u>r</u>egulator of <u>a</u>cidification of <u>v</u>acuoles and <u>e</u>ndosomes), consists of Rav1p, Rav2p, and Skp1p. Mammalian Skp1 has not been detected previously as part of Rabconnectin-3 and is not enriched in the sample. Interestingly, the t-SNARES syntaxin-1A/B and SNAP25 were detected as enriched in the vesicle preparation (**Fig. 1B**, *blue*). The presence of these proteins suggests that some fraction of the sample is derived from the presynaptic membrane or from SVs that were interacting with the presynaptic membrane.

### SVs isolated via SidK are capable of ATP-driven acidification

The SV preparation demonstrates robust ATP-dependent acidification (**Fig. 1C**, *light blue curve*) even before removal of V_1_ complexes by gel filtration chromatography. In this assay, SVs are incubated with the fluorophore 9-amino-6-chloro-2-methoxyacridine (ACMA) and fluorescence recorded. Addition of ATP (**Fig. 1C**, *black arrow*) leads to proton pumping into the SV lumen, which traps and accumulates ACMA, quenching its fluorescence. Adding the K^+^/H^+^ antiporter nigericin (**Fig. 1C**, *blue arrow*) allows rapid exchange of protons from the vesicle lumen with K^+^ ions from the assay buffer, neutralizing the lumen and leading to fluorescence recovery. In contrast, when vesicles are acidified by the addition of ATP and the V-ATPase-specific inhibitor bafilomycin A1 added instead of nigericin (**Fig. 1C**, *red curve*), fluorescence recovers slowly, consistent with protons gradually leaking from the vesicle lumen. SVs preincubated with bafilomycin A1 do not undergo ATP-driven acidification and nigericin-induced neutralization (**Fig. 1C**, *dark blue curve*). SVs separated from free V_1_ complexes by gel filtration chromatography display the same ATP-driven acidification (**Fig. S1C**), although a substantially larger volume of vesicle solution is required for this assay, indicating loss of SVs during gel filtration.

### Imaging of SVs by cryo-EM

We next imaged the SV preparation by cryo-EM on custom nanofabricated holey gold grids^22^ coated with a layer of graphene oxide^23^. Micrographs (**Fig. 1D**) and electron tomograms (**Movie S1**) show vesicles with a diameter of ∼40 nm, in agreement with previous measurements of SVs^13^. The tomograms show the SVs are mostly intact, although they appear to have a flattened surface where they contact the graphene oxide film (**Movie S1**). Both tomograms and 2D micrographs reveal multiple densities protruding from the vesicle membranes, consistent with the V_1_ region of the ∼20 nm long V-ATPase complex (**Fig. 1D**, *red arrows*). Previous mass spectrometry quantification of the V_1_ subunit B in a SV preparation estimated there was on average a single V-ATPase complex per vesicle^24^. However, the images obtained here reveal as many as seven V_1_ complexes protruding from the vesicle membrane (e.g. **Fig. S1B**, *red arrows*). There are several possible explanations for this discrepancy. First, our negative stain EM before gel filtration chromatography shows that detached V_1_ complexes are abundant in homogenized brain tissue, suggesting that dissociated V-ATPases could be more common that assembled V-ATPases. With V_1_ complex dissociated from the SV, quantification of subunit B from the V_1_ region would lead to an underestimate of the number of V-ATPase per vesicle. Consistent with V_1_ dissociating from SVs, recent electron microscopy of SVs isolated using a recombinant antibody that binds the vesicular GABA transporter noted that V_1_-like densities could be detected only in some vesicles^25^. A second reason for the discrepancy could be that our procedure, which relies on the affinity of SidK for V_1_, may preferentially purify SVs with multiple V-ATPases. Therefore, while mass spectrometry quantification of subunit B could underestimate the average number of V-ATPases per SV owing to loss of V_1_, our preparation may overestimate this quantity owing to enrichment of vesicles with an unusually large number of V-ATPase complexes.

### Three-dimensional structure of the V-ATPase within the native SV membrane

Images of V-ATPase protruding from SV membranes were selected for computational image analysis and used with single particle methods to calculate 3D maps of the enzyme (**Fig. 2A**, **Fig. S2**, **Table S1**). Three-dimensional image classification allowed for separation of the dataset into classes corresponding to the three main rotational conformations of the enzyme, known as Rotational States 1, 2, and 3 ^6,26^. The nominal overall resolutions of the maps were 3.9 Å for State 1 (∼25% of particle images), 3.6 Å for State 2 (∼29% of particle images), and 3.5 Å for State 3 (∼46% of particle images). Local resolution ranged from 7.2 to 3.4 Å for the State 3 map. The 3D maps reveal the V-ATPase protruding from a spherical cap of lipid bilayer with a radius consistent with the ∼40 nm diameter of SVs. Local refinement of the V_1_ and V_O_ regions improved their overall resolutions to 3.2 and 3.8 Å, respectively, with local resolutions ranging from 5.2 to 2.4 Å and 4.6 to 2.4 Å, respectively. Therefore, despite images being obtained in the protein-rich native lipid bilayer of a functional synaptic vesicle, the maps are of sufficient resolution for straightforward construction of an atomic model of V-ATPase (**Fig. 2B** and **C**).

**Figure 2.**
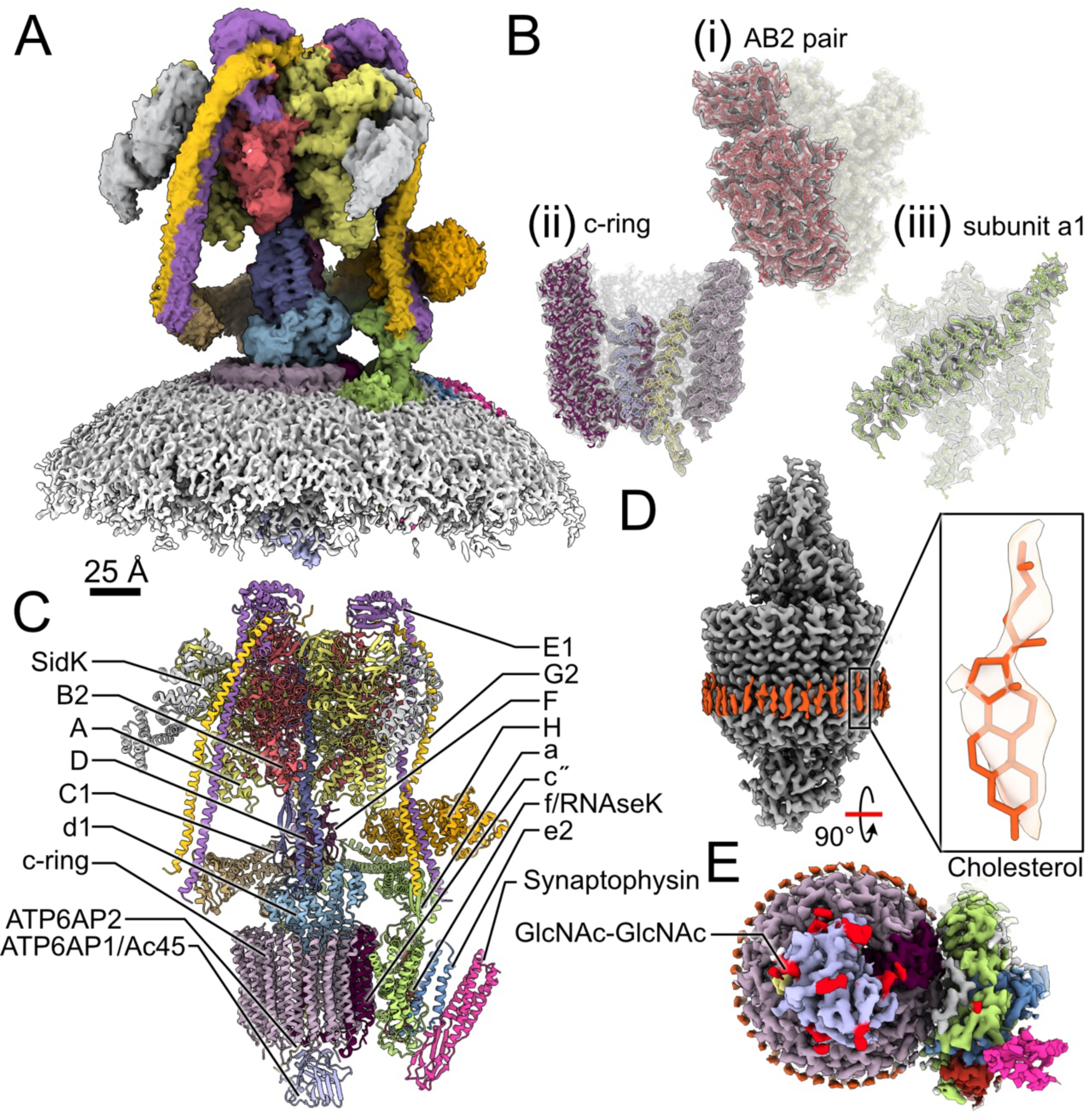
Structure of V-ATPase in the native SV membrane. **A,** Structure of V-ATPase in a spherical cap of lipid bilayer from the SV membrane. **B,** Examples of high-resolution map density with an atomic model for a pair of A and B2 subunits in the V_1_ region (**i**) part of the c-ring in the VO region (**ii**), and the C-terminal domain of subunit a1 in the V_O_ region (**iii**). **C,** Complete atomic model for the synaptic vesicle V-ATPase. **D,** Densities corresponding to cholesterol from the inner leaflet of the SV membrane decorate the c-ring. **E,** Densities corresponding to the carbohydrates GlcNAc or GlcNAc-GlcNAc are attached to subunit ATP6AP1/Ac45 and subunit a1.

### Sterols and carbohydrates decorate the V-ATPase membrane region

In the map, the proton conducting c-ring of the V-ATPase is decorated with regularly spaced densities within the luminal leaflet of the lipid bilayer (**Fig. 2D**, *orange densities*). These densities have the characteristic size and shape of cholesterol. SVs are cholesterol rich, with the lipid reported to make up between 33 and 40% of the membrane^27^. Consequently, the regularly spaced densities were modelled as cholesterol. Sterols are necessary for the function of yeast V-ATPase^28^ and extraction of cholesterol from SV membranes with methyl-β-cyclodextrin inhibits their ATPase activity^29^. The cholesterol molecules in the structure are similar to densities seen bound to the detergent-solubilized yeast V-ATPase c-ring^30^. However, the yeast V-ATPase structure was determined using the sterol-derived detergent glyco-diosgenin, making it impossible to distinguish between glyco-diosgenin and native sterols. In contrast, the present structure was determined without detergents, indicating that bound cholesterol is a bona fide feature of the V-ATPases structure. Remarkably, the position of these cholesterols would prevent rotation of the c-ring owing to steric clashes with subunit a. Therefore, the structure suggests that during ATP-driven proton pumping cholesterol molecules continuously bind and unbind from the rotating c-ring. This model resembles the bound cardiolipin in the rotor ring of mitochondrial ATP synthase, where the specialized lipid has been proposed to lubricate rotation of the rotor in the membrane^31^.

Fitting of the atomic models for subunit ATP6AP1/Ac45, subunit a1, and subunit e2 into the corresponding regions of the map revealed additional density that is not occupied by the protein (**Fig. 2E**, *red densities*). This extra density is found near Asn residues 255, 267, 290, 297, 344, 351, and 399 of subunit ATP6AP1/Ac45; residue Asn489 of subunit a1; and residue Asn70 of subunit e2. Corresponding residues have previously been found to be glycosylated in the human V-ATPase and the first two sugar units were modelled as two consecutive N-acetylglucosamine (GlcNAc)^15^. The structure shown here reveals that even in the context of a native SV, the polysaccharides are not ordered past their second sugar moiety.

### V-ATPase interacts with synaptophysin in synaptic vesicles

The cryo-EM map reveals a protein density in the SV membrane attached to the V-ATPase that cannot be explained by the known subunits of the enzyme (**Fig. 3A**, *pink*). Despite extensive 3D classification, it was not possible to obtain a subpopulation of particle images that lack this density. This observation, combined with the high density for the additional protein in the map, indicate that the additional protein is a stoichiometric component of V-ATPase in SVs. Nonetheless, the density for this protein had quite poor resolution except where it contacts the rest of the V-ATPase, likely owing to variable position around where it is bound to the enzyme. The density shows four transmembrane α-helices and a small mixed α/β domain in the SV lumen (**Fig. 3B**). This fold resembles the predicted structures for MARVEL-domain proteins that are resident in SVs: synaptophysin, synaptoporin, pantophysin (also known as synaptophysin-like protein 1), mitsugumin (also known as synaptophysin-like protein 2), and synaptogyrin. Of these proteins, synaptophysin, synaptoporin, synaptogyrin-1, and synaptogyrin-3 were detected in the SV preparation (**Fig 1B**, *orange*, **Supplementary Data 1**). The AlphaFold models for both synaptophysin and synaptoporin^32^ fit the overall shape of the density with high fidelity (**Fig. 3B**), whereas synaptogyrin-1 and −3 lack the mixed α/β luminal domain. This region of the protein, where it contacts V-ATPase, is well-resolved with clear density for amino acid side chains (**Fig. 3C**). The side chain densities indicate bulky residues that align with Tyr75 and Phe90 of the fitted synaptophysin AlphaFold model but not with the corresponding Thr55 and Ala70 residues of the synaptoporin model (**Fig. 3D**). Therefore, we conclude that this additional protein is synaptophysin. Nonetheless, owing to the image averaging inherent in single particle cryo-EM, we cannot exclude the possibility that some of the V-ATPase complexes are bound to a different MARVEL-domain protein. The interaction between V-ATPase and synaptophysin involves His72, Gln73, Tyr75, Lys88, Phe90, and Asp94 from synaptophysin; Arg482 and Glu492 from the proton-conducting subunit a1; Arg77, Phe78, Leu79, and Glu81 from subunit e2; and Pro55 and Tyr59 from subunit f (**Fig. 3A**, *close-up*). Interestingly, these interactions are the only known structural role for V-ATPase subunits e2 and f. Both synaptophysin and synaptobrevin-2 were previously reported to interact with the V_O_ complex^33^. However, we could not detect density for synaptobrevin-2 adjacent to the V-ATPase in the present structure. More recently, cross-linking mass spectrometry detected interactions between synaptophysin and both subunit d1 and the soluble N-terminal domain of subunit a1^34^.

**Figure 3.**
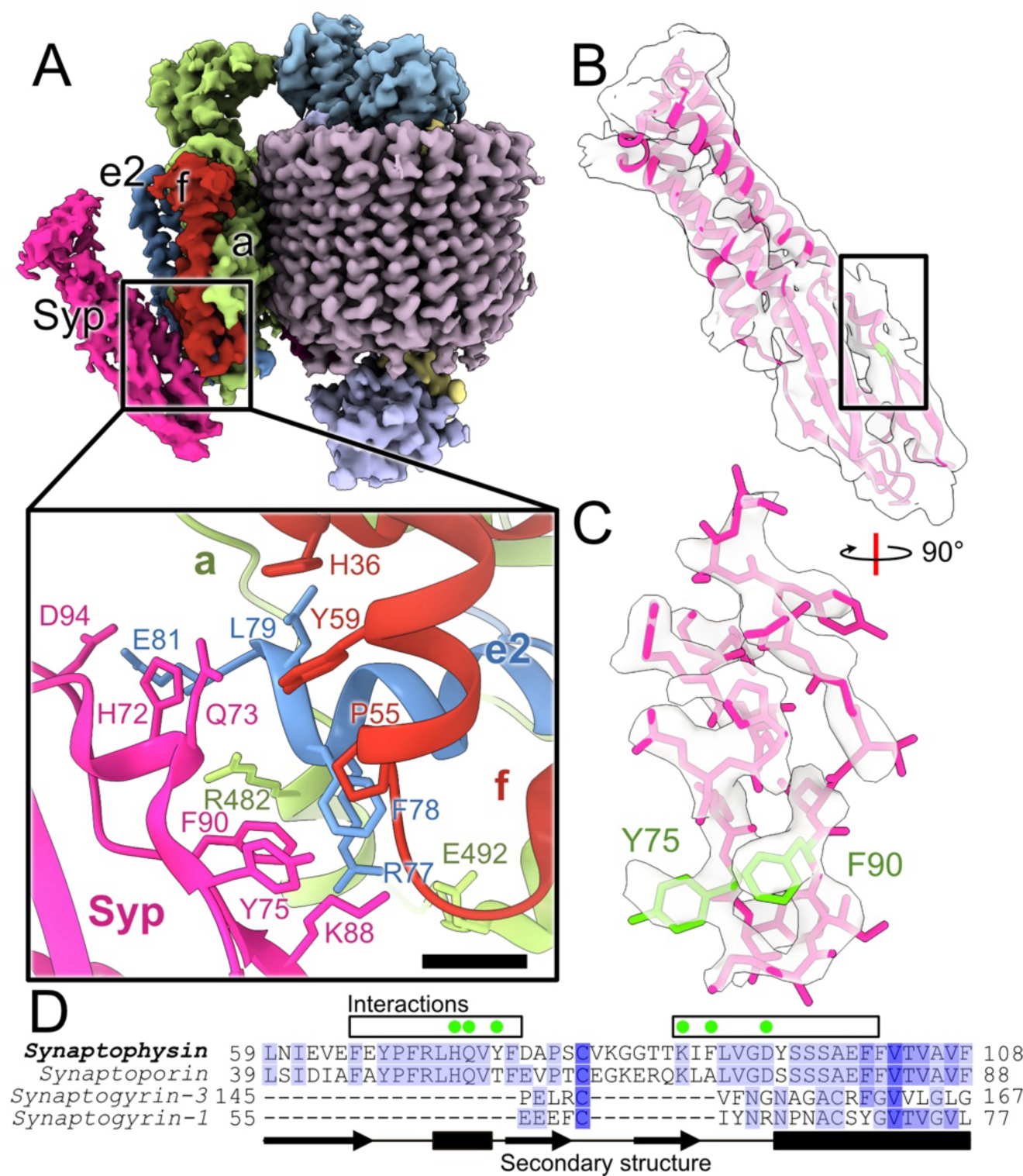
Synaptophysin is a subunit of the V-ATPase in SVs. **A,** A close-up view of the V_O_ region showing density from synaptophysin attached to subunits a, e2, and f. Close-up: amino acid residues involved in the interaction. Scale bar, 5 Å. **B,** The additional density fits the MARVEL domain proteins synaptophysin and synaptoporin. **C,** Synaptophysin, but not synaptoporin, possesses bulky side chains that are consistent with high-resolution density where the MARVEL domain protein contacts the rest of V_O_. **D,** Multiple sequence alignment for synaptopysin, synaptoporin, and synaptogyrin-1 and −3. Conserved residues are shaded in blue. Empty boxes indicate the well-resolved region of the protein and green circles indicate residues involved in the V_O_ region interaction. Predicted secondary structure elements are indicated (black arrows and filled rectangles).

Synaptophysin is the most abundant SV protein by weight, with ∼32 copies per vesicle accounting for ∼10% of the total SV protein content^24^. Owing to its abundance in SVs, synaptophysin is often used as a marker for neuroendocrine tumours. The protein can form a hexameric homo-oligomer^35,36^ and interacts with the v-SNARE synaptobrevin-2 ^37,38^, chaperoning the assembly of the v- and t-SNAREs between the SV and the presynaptic membrane^39^. Surprisingly given its evolutionary conservation and abundance, synaptophysin knock-out mice reproduce normally and form morphologically normal SVs that can release neurotransmitters^40,41^. However, neurons from knockout mice show decreased SV endocytosis both during and after sustained neuronal activity^42^ and inefficient retrieval of synaptobrevin-2 from the plasma membrane^43,44^. Human mutations in synaptophysin are linked to intellectual disability^45^ and knockout mice have learning deficits^46^. Nonetheless, this phenotype is relatively subtle for a conserved and abundant SV protein, a paradox that has been suggested to result from other proteins compensating for synaptophysin’s absence or because defective neurotransmission occurs only when neurons fire at high frequency^39,47^. The stoichiometric interaction of V-ATPase with synaptophysin in the SV membrane suggests that, in addition to synaptobrevin-2, synaptophysin may help recover V-ATPase to the SV membrane following neurotransmitter release. The angle between synaptophysin and the rest of V_O_ matches the highly curved SV membrane (**Fig. 2A** and **3A**) but the observed flexibility around the interaction site between synaptophysin and V_O_ suggests that in a planar lipid bilayer synaptophysin could pivot closer to the enzyme to form additional contacts.

### SV loading conditions induce partial V-ATPase dissociation

The ability to image SVs by cryo-EM presents a unique opportunity to understand the consequences of SV acidification and loading on V-ATPase structure. We first tested the ability of our SV preparation to load with glutamate by mixing vesicles with different concentrations of glutamate (0 to 15 mM) prior to adding ATP in acidification assays. Incubating with glutamate resulted in diminished fluorescence quenching (**Fig. 4A**), consistent with VGLUT consuming some of the proton motive force (Δ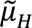+) established by V-ATPase to produce a glutamate gradient across the SV membrane^48^. To determine whether SV loading causes changes in the structure of V-ATPase, we next incubated SVs in a buffer with 2 mM ATP and 16 mM glutamate, mimicking conditions used in the acidification assay before freezing the vesicles for cryo-EM (**Fig. 4B**). Initial image analysis suggested fewer intact V-ATPase complexes than we observed with the non-treated SV sample. This loss of V_1_ complex is unlikely to be the result of a pH change as reagents were buffered and their pH measured.

**Figure 4.**
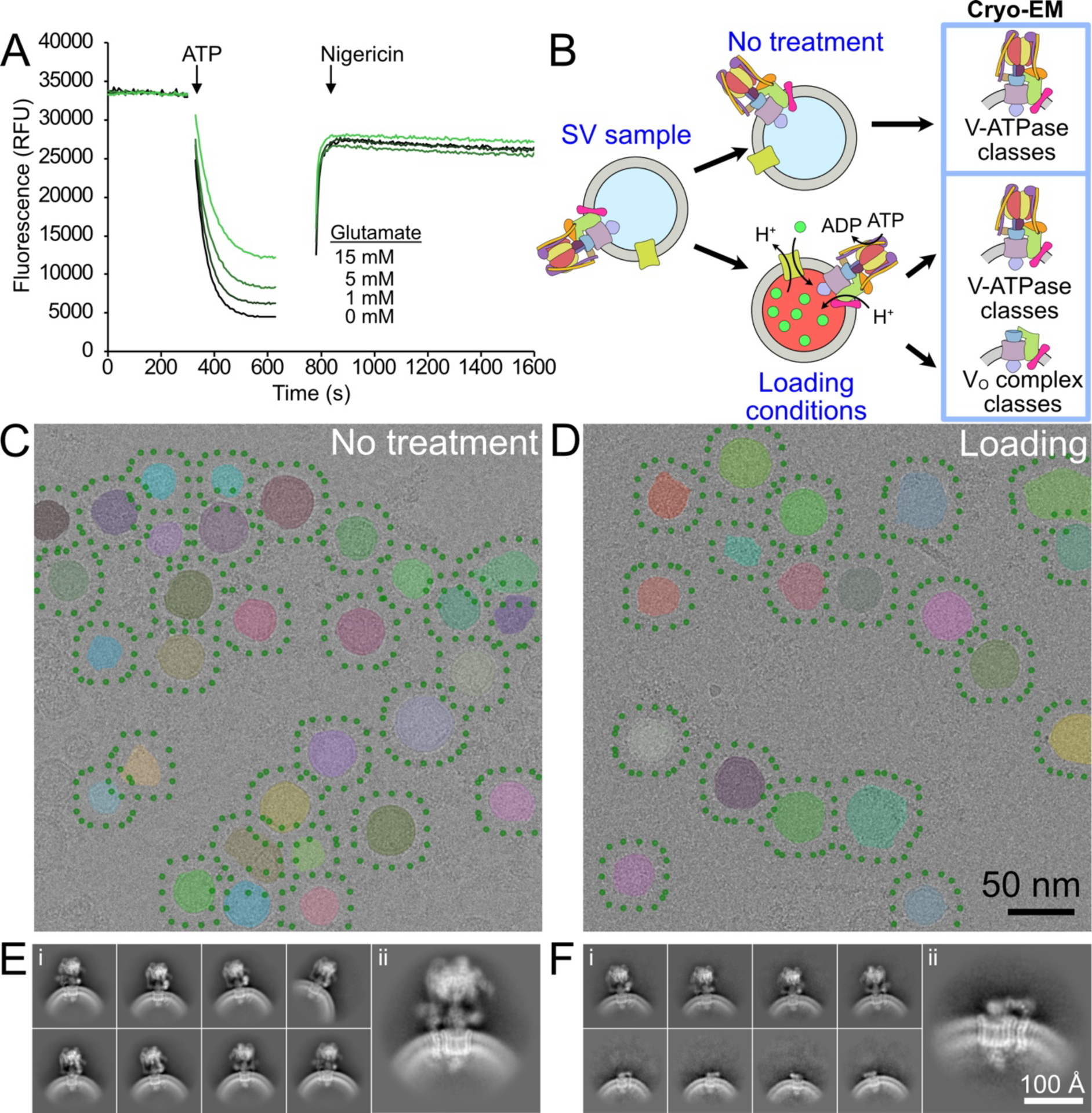
Loading of synaptic vesicles induces V-ATPase dissociation. **A,** Incubation of SVs with increasing concentrations of glutamate leads to decreased vesicle acidification. **B,** Cryo-EM of SVs treated in conditions that drive loading produces class averages of both V-ATPase and dissociated V_O_ complexes. **C,** Automated selection of vesicles (semitransparent shapes) and exhaustive selection of possible V-ATPase locations (green dots) without treatment. **D,** Automated selection of vesicles (semitransparent shapes) and exhaustive selection of possible V-ATPase locations after treatment with SV loading buffer containing 1 mM ATP and 8 mM glutamate (green dots). **E,** 2D classification of exhaustively-selected particle locations without treatment with SV loading buffer. **F,** 2D classification of exhaustively-selected particle locations after treatment with SV loading buffer.

To further characterize the SVs treated with loading buffer, we used a newly developed computer program that allows automated and unbiased detection of vesicles in images and exhaustive selection of the possible particle locations around the bounding lipid bilayer (see methods). We applied this tool to micrographs of SVs without (**Fig. 4C**) and with (**Fig. 4D**) treatment in loading conditions. The exhaustively selected images of regions of lipid bilayer were subjected to 2D classification, and classes showing V-ATPase were selected and separated further (**Fig. S3**). Following this process, as expected, classes showing intact V-ATPase could be produced from vesicles that had not been treated in loading conditions (**Fig. 4E**). However, after loading, 2D classes appeared that showed not just intact V-ATPase (**Fig. 4F**, *top row*), but also isolated V_O_ complexes (**Fig. 4F**, *bottom row*), indicating that the loading conditions are sufficient to induce dissociation of V-ATPase. Indeed, nearly ∼37% of the particle images showed V_O_ complexes rather than intact V-ATPase complexes after loading. In contrast, classes showing V_O_ complexes could not be identified from images of untreated SVs. Using particles from the classes to train a machine-learning particle selection algorithm^49^ led to 2D classes with high-resolution features for both intact V-ATPase (**Fig. 4E**, *enlarged image*) and V_O_ complex (**Fig. 4F**, *enlarged image*).

## Discussion

Single particle cryo-EM of membrane proteins typically involves extracting the protein from the lipid bilayer with detergent and determining its structure in detergent micelles or following reconstitution into another membrane mimetic. However, previous structures of the mammalian brain V-ATPase in detergent did not provide any indication that synaptophysin interacts with the enzyme^6,10^. This interaction was only resolved by determining the structure in the native SV membrane without using detergents or other reagents that perturb the lipid bilayer. Previous work used cryo-EM to image membrane proteins reconstituted into proteoliposomes^50–52^ and even in vesicles derived from the cells used for heterologous overexpression^53^. While these earlier experiments enable examination of the effects of transmembrane voltage and ion gradients on protein structure, they would not have allowed for identification of the interaction of V-ATPase with synaptophysin. Tomograms like the one in Supplementary Movie 1 would likely facilitate selection of more V-ATPase complexes per vesicle, and sub-tomogram averaging would probably yield a high-resolution structure of the V-ATPase:synaptophysin complex. However, the approach employed here allowed use of efficient untilted image data collection and optimized single particle cryo-EM computational methods, which outweighed the advantage of electron tomography.

Fluorescence microscopy of individual SVs fused with pH-sensitive fluorescent large unilamellar vesicles (LUVs) revealed that V-ATPase undergoes slow mode switching between functional states^54^. The first mode is an active state where the SV-LUV fusion acidifies, the second is an inactive state where vesicles temporarily leak protons through the membrane, while in the third state the fused SV-LUVs abruptly neutralize when the V-ATPase itself enables proton leak. It is unlikely that the dissociation of V_1_ from V_O_ seen in the present experiment explains the temporary inactive state, because both the cryo-EM and fluorescence microscopy experiments lacked the cellular machinery needed for V-ATPase reassembly. However, the full dissociation of V-ATPase seen by cryo-EM could explain the proton-leak state. Although dissociated yeast V_O_ complex is impermeable to protons^55^, the same property has not been demonstrated for the mammalian enzyme.

Previous work has shown that loading of chromaffin granules in PC12 cells leads to dissociation of V_1_ from V_O_ ^3^. Further, dissociation of the V-ATPase following synaptic vesicle loading is required for neurotransmitter release from hippocampal neurons^2^. While Rabconnectin-3 is needed to reassemble V_1_ with V_O_, it is not known if dissociation of V-ATPase requires additional proteins or is an intrinsic property of the enzyme in SVs. The experiments described here indicate that V_1_ readily dissociates from V-ATPase in SVs in conditions that induce SV loading, highlighting the labile nature of the complex and suggesting that loading alone is sufficient to induce dissociation. Spontaneous dissociation implies that control of whether V-ATPase is assembled and active or dissociated and inactive can be achieved simply by regulating its reassembly by Rabconnectin-3 (**Fig. 1A**). This finding is consistent with previous results that suggest that ATP hydrolysis leads to V-ATPase dissociation in yeast cells^56^ and with the detergent-solubilized enzyme from *Manduca sexta* midgut^57^. Interestingly, despite the loss of V_1_ from the SV during proton pumping, ACMA fluorescence quenching assays do not show gradual neutralization of the SV during the assay (e.g. **Fig. 1C** and **4A**). This observation is consistent with the H^+^:ATP ratio of the V-ATPase setting the ultimate Δ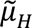+ across the SV membrane^6,26^, and the number of pumps determining the rate at which that proton motive force can be achieved. Dissociation of V_1_ from the SVs could mark vesicles that are loaded and ready for release^2^.

## Methods

### Purification of SidK

SidK_(residue 1-278)_ was purified as described previously^6^. Briefly, competent CodonPlus *Escherichia coli* cells were transformed with pET-28a-His_6_-TEV-SidK_(1-278)_-3×FLAG. Individual colonies were inoculated into 5-10 mL Luria Broth (LB) supplemented with 50 μg/mL kanamycin and grown at 37°C overnight. LB (1 L) supplemented with 0.4 % (w/v) D-glucose and 50 μg/mL kanamycin was inoculated with 5 mL overnight culture and grown at 37°C with shaking at 200 rpm. Once an OD600 of ∼0.7 was reached, protein expression was induced with addition of 1 mM isopropyl β-D-1-thiogalactopyranoside (IPTG) and temperature was lowered to 16°C overnight. Cells were collected at 5,250 ×g for 45 min at 4°C and used immediately or stored at –80°C.

Cell pellets were thawed and resuspended in 40 mL HisTrap buffer A (50 mM Tris-HCl pH 7.4, 300 mM NaCl, 25 mM imidazole) per 1 L cell culture. Cells were lysed by sonication (Misonix Sonicator 3000; total on time 3 min with 5 s on, 10 s off, power level 6.5). Cell debris and insoluble protein was removed by centrifugation at 38,000 ×g for 30 min at 4°C. The pH of the supernatant was adjusted to 7-8 before 5 min incubation with 0.1 mg/mL DNAse I and filtered with a 0.45 μm syringe filter. Filtered supernatant was loaded onto a HisTrap HP column (Cytiva) equilibrated in HisTrap buffer A, before washing with six column volumes of HisTrap buffer A and elution in HisTrap buffer B (50 mM Tris-HCl pH 7.4, 300 mM NaCl, 300 mM imidazole). Eluted fractions were pooled and TEV protease was added to cleave the N-terminal His tag. The sample was dialysed with TEV overnight against 2 L Tris-buffered saline (TBS; 50 mM Tris-HCl pH 7.4, 150 mM NaCl) with 1 mM dithiothreitol (DTT). DTT was removed by further dialyses against TBS without DTT, first for 45 min and then for 2 h before being filtered with a 0.45 μm syringe filter and loaded onto a HisTrap column equilibrated in TBS. The column was washed with TBS, followed by HisTrap buffer A. SidK-containing fractions were pooled, buffer-exchanged to TBS, and concentrated with an Amicon Ultra 10 kDa MWCO concentrator (Millipore). Protein aliquots were frozen with liquid nitrogen and stored at −80°C.

### Isolation of synaptic vesicles

A protocol for isolating synaptic vesicles from rat brain using SidK affinity to V-ATPase was developed by combining aspects of protocols for purification of synaptic vesicles^20^ and purification of detergent solubilized rat brain V-ATPase^6^. All steps were performed on ice or at 4°C. SidK_(1-278)_-3×FLAG (∼0.85 mg) was bound to 1.5 mL of M2-FLAG affinity gel resin (Sigma) and flow through was recirculated onto the column 6-7 times. The column was then washed with 5-10 column volumes of TBS. The brain of a ∼3-12 months old Sprague Dawley rat (*Rattus norvegicus*) was washed three times with 10 mL of ice-cold KPBS (10 mM KH_2_PO_4_ pH 7.25, 136 mM KCl). The brain was homogenised in 5 mL KPBS with 30 strokes of a tight-fitting glass-Teflon pestle with scores at 900 rpm using a Caframo 2002 Compact Digital homogeniser. Sample was divided between eight 1.5 mL Eppendorf tubes and centrifuged at 17,000 ×g for 3 min to remove unbroken cells and large debris. Supernatant was filtered through 0.22 µm PES Millex-GP (Millipore Express) syringe filters, loaded onto the M2-FLAG column with immobilised SidK, and recirculated 4-6 times. The column was washed with 15 mL TBS and eluted with 3 mL of 150 µg/mL 3×FLAG peptide (Genscript) in TBS. The sample was concentrated in a 4 mL Amicon Ultra 100 kDa MWCO concentrator (Millipore) and then injected onto a Superose 6 10/300 column (Cytiva) on an AKTA Pure (Cytiva), previously equilibrated in TBS. Vesicle-containing fractions were pooled and concentrated with a 0.5 mL Vivaspin 100 kDa MWCO PES concentrator (Sartorius) to ∼20-40 µL for cryo-EM sample preparation.

### Assay of synaptic vesicle acidification

An assay for acidification of synaptic vesicles was developed by combining aspects of an earlier synaptic vesicle assay^58^ with a related assay for bacterial inverted membrane vesicles^59^. SVs were diluted in KCl Assay Buffer (20 mM HEPES-KOH pH 7.4, 150 mM KCl, 3 µM 9-amino-6-chloro-2-methoxyacridine [ACMA], 2 mM MgSO_4_) in 96-well microwell plates. ACMA fluorescence was monitor with excitation at 410 nm and emission at 490 nm in a Synergy Neo2 HTS plate reader held at 22°C. After ∼5 min, reactions were initiated by addition of ATP solution that had been adjusted to pH 7.4 with NaOH to reach a final concentration of 1.6 mM. Proton pumping was allowed to proceed for ∼20 min before neutralizing the vesicles by adding 1 µL of 100 µM nigericin in water. Where added, glutamate stock solution was adjusted to pH of 7.4 before use.

### Sample preparation for mass spectrometry

Samples of purified SVs were separated on an SDS-PAGE gel, along with samples of homogenized rat brain at different dilutions following the 17,000 ×g for 3 min centrifugation step. Gels were stained with Coomassie. The dilution of rat brain lysate where staining was comparable to the SV lane was selected as the “input” for experiments. Each gel lane was cut into small pieces and separated into four tubes. The gel pieces were washed two times with 150 µL of 100 mM ammonium bicarbonate (ABC) on ice. The gel fragments were then incubated with 150 µL of dehydrating solution (95% ethanol and ABC mixed 19:1) on ice for 20 min to dehydrate the gel. Dehydrating solution was removed, the gel fragment washed with ABC, and another 100 µL of the dehydrating solution added. After removal of the dehydrating solution, 80 µL of 5 mM DTT in ABC was added to each tube and the tubes were incubated at 55°C for 30 min. The gel fragments were next treated with 100 µL of dehydrating solution for 10 min and then 80 µL of 50 mM iodoacetamide in ABC for 45 min at room temperature in the dark. The iodoacetamide solution was removed and dehydrating solution (100 µL) was added to each tube for 10 min. For trypsinolysis, 840 ng trypsin (Promega) in 30 μL ABC was added to each tube, incubated for 10 min at room temperature, and then an additional 40 µL of ABC was added to each tube before allowing the sample to digest at 37°C overnight. For digestion with chymotrypsin (Promega), the same process was followed but digestion was allowed to proceed at 25°C overnight. After overnight incubation the reaction was mixed and briefly centrifuged, the solution removed and added to a fresh tube, and 50 µL of 5% formic acid was added to the gel fragments. The formic acid solution was incubated with the gel fragments for 20 min on ice, mixed and centrifuged, and this solution added to the solution from the overnight incubation. The addition of formic acid solution and collection of formic acid extracts was repeated, and the collected peptide solutions were dried by Speedvac with no heating and placed on dry ice before storing at −80°C.

### Liquid chromatography-mass spectrometry

For LC-MS/MS, peptides isolated from gel fragments of the same lane were combined into a single sample that was analyzed with nano high-performance liquid chromatography coupled to mass spectrometry. One-twelfth of the sample was loaded onto an EvoSep Evotip Pure (EV2018) tip following the manufacturer’s instructions. Peptides were separated on an EvoSep Performance (EV1109) column (8 cm × 150 µm with 1.5 µm C18 beads) held at 40°C with the 60SPD pre-formed acetonitrile gradient (∼20 min total gradient time) generated by an Evosep One system, and analyzed on a Bruker timsTOF Pro 2 mass spectrometer. The Evosep was coupled to the mass spectrometer using a 20 µm internal diameter electrospray emitter tip (Bruker p/n 1865710). Sample-to-sample injection time was approximately 22 min. Spectra were collected using data-dependent PASEF acquisition and an active exclusion time of 0.4 min, target intensity of 17,500, and an intensity threshold of 1,750. Trapped ion mobility was operated with fixed accumulation and ramp times of 100 ms across an inverse mobility (1/K_0_) range from 0.85 to 1.3 V·s/cm^2^, with four mobility ramps across the PASEF cycle. The cycle time was 0.53 s. The MS1 scan had a mass range of 100-1700 m/z. Singly-charged ions were excluded from MS2 fragmentation with a mobility-selective polygonal filter and auto calibration was off.

### Mass spectrometry database analysis

Data were searched using MSFragger 3.7 within the ProHits LIMS. The FASTA database is from UP000002494 rat Uniprot, with one protein per gene. The database included additional contaminant proteins^60^ as well as ten biotin ligase enzymes. Cysteine carbamidomethylation (57.0214) was set as a fixed modification. Acetylated protein N-term and oxidated methionine were set as variable modifications. Precursor mass tolerance was set to 40 ppm on either side. Fragment mass tolerance was set to 40 ppm. Enzymatic cleavage was set to either trypsin or chymotrypsin with two missed cleavages. MSBooster and Percolator were turned on. PSM Validation cmd line: -minprob 0.05 --decoyprobs --ppm --accmass --decoy DECOY –nonparam --expectscore. Percolator cmd line: --only-psms --no-terminate --post-processing-tdc. PTMprophet cmd line: KEEPOLD STATIC EM=1 NIONS=b MINPROB=0.5. ProteinProphet cmd line: --maxppmdiff 30. FDR filter and report cmd line: --sequential --razor --prot 0.01 -- picked --mapmods. All other parameters were default.

### Negative stain electron microscopy

Maxtaform copper/rhodium 400 mesh electron microscope grids (Electron Microscopy Sciences) were coated with collodion film and a thin layer of carbon deposited by evaporation of carbon in an EM ACE200 sputter coater (Leica). Grids were rendered hydrophilic by glow discharge in air with a Pelco easiGlow glow discharge device for 15 s. SV sample (∼3 µL) was placed on the grid and SVs were allowed to adsorb for 2 min. The sample was washed off with three drops of water and stained with 2% uranyl acetate before being allowed to dry in air. Images were obtained with a TF20 electron microscope (FEI Company) operating at 200 kV and equipped with a Gatan Orius CCD camera at 25,000× magnification, which corresponds to a calibrated pixel size of 2.61 Å/pixel.

### Graphene oxide grid preparation

Graphene oxide-coated grids were prepared on holey gold grids as described^23^. Briefly, 60 µL graphene oxide stock at ∼2 mg/mL concentration, was diluted with 740 µL of 5:1 methanol:MilliQ water and agitated by stirring to remove chunks. In-house nanofabricated holey gold-coated grids^22^ were submerged in MilliQ water. Graphene oxide was spread on the surface of the water by adding 230 µL of the suspension onto the surface of the water at 50 µL/min using a syringe pump. A peristaltic pump was used to slowly removed the water and lower the graphene oxide layer onto the gold grids. Grids were dried and screened with a TF20 electron microscope operating at 200 kV to confirm the presence of a single layer of graphene oxide.

### CryoEM data collection

Sample (3 µL) was added to the graphene oxide-coated gold grids, previously glow discharged in air for 15 s, and allowed to adsorb for 1 min in the blotting chamber of a TFS Vitrobot Mark IV at 4°Ç 95–100% humidity. Samples were blotted for 3 s with blot force 5 and frozen by plunging into liquid ethane. To mimic loading conditions obtained in acidification assays, ATP was added to 2 mM and glutamate to 16 mM by mixing SVs with a buffered stock solution (50 mM Tris, 150 mM NaCl, 8 mM ATP, 16 mM MgSO_4_, 64 mM glutamate, 48 µM CaCl_2_, pH adjusted to 7.4 with NaOH) with half the concentration added before concentrating the sample (total time 1 h) and half added immediately before grid freezing.

For single particle analysis, samples were imaged with a ThermoFisher Scientific Titan Krios G3 electron microscope, operating at 300 kV and equipped with a Falcon 4i camera. Data was acquired with TFS EPU software v3.7. Movies were acquired in electron event representation mode^61^ and subsequently divided into 30 exposure fractions. The exposure rate used with the camera was 9.6 electron/pixel/s and a total exposure of ∼37.5 electrons/Å^2^ was applied. The calibrated pixel size was 1.03 Å. Electron tomograms were collected with TFS Tomography v5.16. Tilt series of ±60° were collected with 2.5° steps. The calibrated pixel size was 2.2 Å and an exposure of ∼6.5 electrons/Å^2^ was used for each of the 24 exposures. Tomograms were calculated with AreTomo^62^. Tomograms were rescaled, their orientation with respect to the 3D volume axes adjusted, and movies rendered with Fiji^63^.

### Single particle cryo-EM image analysis

Data were processed with cryoSPARC v4.4.0^64^. Movies were aligned with the patch motion correction job and contrast transfer function (CTF) parameters were estimated initially with the patch CTF job. Particle images (247) centred on the V_1_ region of V-ATPase were selected manually from a subset of 32 micrographs. 2D classification yielded classes that showed the V-ATPase complex, and images from these classes were used as templates for particle selection from a set of 715 micrographs. Particle images from these micrographs that contributed to 2D classes with visible V-ATPase complex were used to train a Topaz model^49^ that allowed extraction of particle images from 1,721 micrographs. These images were subjected to 2D classification and particles contributing to classes with V-ATPase were used in *ab initio* reconstruction with three classes, yielding a single 3D map showing V-ATPase in the lipid bilayer. This 3D map was refined with heterogeneous and homogeneous refinement.

Templates were generated from this first 3D map and used with the template picker job to locate particle images with views away from the membrane bilayer, including top views. Topaz model training and particle image extraction was iterated with 2D and 3D classification for the different views of the V-ATPase, to increase particle image numbers and improve centering. Particle images were pooled to create a set of 1,060,637 images and re-extracted from micrographs in 512×512 pixel boxes. Rounds of *ab initio* and heterogeneous refinement generated a final set of 432,463 particle images. Heterogeneous refinement was performed with copies of the best V-ATPase map to generate maps of the three rotational states, which were refined with non-uniform refinement^65^ to give maps of Rotational State 1 (108,773 particle images, 3.9 Å resolution), Rotational State 2 (123,220 particle images, 3.6 Å resolution), and Rotational State 3 (198,766 particle images, 3.5 Å resolution).

Reference-based motion correction was applied to the State 3 particle image stack and particles were re-extracted in 300×300 pixel boxes, providing 198,533 particle images, centering on different regions of the map for focused refinement. Particle subtraction and local refinement were performed focussing on the V_O_ region, V_1_ region, and the peripheral stalks, yielding maps at 3.2, 3.2, and 3.8 Å resolution, respectively. The resolution of these maps was sufficient to allow for model building of the V-ATPase starting with a model comprising PDB 6VQ8 ^6^ with subunits H and Ac45/PRR from PDB 7UZF. An AlphaFold model for synaptophysin (P07825, residues 11-224) was docked into the corresponding low-resolution density and sidechains were built where density was visible. The overall model was refined iteratively with ISOLDE^66^ and Phenix real-space-refinement (version 1.21-5207)^67^. Interacting residues between synaptophysin and subunits a1, e2, and f of V-ATPase were identified with LigPlot+ v2.2.8 ^68^.

### Computational comparison of micrographs before and after SV loading

SVs were identified in micrographs with a custom algorithm, VesiclePicker (https://github.com/r-karimi/cryoEM_vesicle_segmentation), implemented in Python. Micrographs were transferred to the program, and possible particle locations transferred back to cryoSPARC with the cryosparc-tools library. Briefly, each micrograph was downsampled 4×4 and an edge-preserving bilateral filter implemented in OpenCV applied (https://opencv.org/]. Putative SVs were identified and segmented with the Segment Anything model^69^. These putative SVs were subjected to a minimum roundness cut-off and minimum and maximum area cut-offs to select vesicles for further analysis. The parameters for filtering, mask generation with Segment Anything, roundness cut-off, and area cut-offs were optimized empirically with a small dataset of manually annotated micrographs. To select V-ATPase and V_O_ particles in SV membranes, the segmentation masks corresponding to SVs were dilated and the locations of the dilated mask edges from the downsampled micrographs were exported to cryoSPARC for extraction from micrographs and exhaustive 2D classification. Classes showing features consistent with V-ATPase or V_O_ were selected for further classification. From 17,148 micrographs with SVs without loading buffer treatment, 3,707,176 candidate particle positions were considered, leading to 142,498 particle images included in 2D classes that showed intact V-ATPase, and none contributing to 2D classes showing V_O_ complexes. For the SVs treated with loading buffer, 7,986 micrographs with SVs yielded 714,003 candidate particle positions, with 16,674 particle images (∼63%) contributing to 2D classes showing intact V-ATPase and 9,601 particle images (∼37%) contributing to classes that showed dissociated V_O_ complexes.

## Acknowledgements

We thank Dr. Brendon Searle for advice on mass spectrometry, Prof. Mike Cousin for discussions on synaptophysin, Dr. Zhijie Li for guidance on electron tomography, Dr. Samir Benlekbir for assistance with cryo-EM data collection, and Dr. Lin Mei for preliminary efforts at cryo-EM of synaptic vesicles. CEC and RK were supported by ResTraComp fellowships from The Hospital for Sick Children. JDT was supported by a Postdoctoral Fellowship from the Canadian Institutes of Health Research (CIHR) and YL was supported by a Canada Graduate Scholarship from the CIHR. GMC was supported by a Mary H. Beatty Fellowship from the University of Toronto. JLR, L-YW, and J-YY were supported by the Canada Research Chairs program. Cryo-EM data were collected at the Toronto High-Resolution High-Throughput cryo-EM facility and enzyme assays performed using infrastructure from the Hospital for Sick Children’s Structural and Biophysical Core Facility, both of which are supported by the Canada Foundation for Innovation and Ontario Research Fund. Mass spectrometry data were collected at the Network Biology Collaborative Center at the Lunenfeld-Tanenbaum Research Institute. This research was supported by CIHR grants to JLR (PJT166152), L-YW (PJT15603, PJT156439, PJT191780), and J-YY (PJT482432) and from the Natural Sciences and Engineering Research Council RGPIN-2023-04676 (JLR), RGPIN-2017-06665 (L-YW), and RGPIN-2022-04849 (J-YY).

## Author contributions

CEC developed the final synaptic vesicle purification procedure, isolated synaptic vesicles, performed, assays, prepared cryo-EM specimens, imaged specimens and performed image analysis, and constructed atomic models. RK developed the vesicle selection software and performed the computational comparison of complexes in synaptic vesicles with and without treatment with loading buffer. SAB developed the initial synaptic vesicle purification procedure and acidification assay procedure. YL, GMC, and JDT contributed to the development of the specimen preparation and image analysis methods for vesicles. CW helped design, performed, and analyzed mass spectrometry experiments. RS helped with functional characterization of the synaptic vesicles. J-YY guided design of mass spectrometry experiments and their analysis. L-YW guided interpretation of findings related to neurobiology. JLR conceived the project, supervised experiments, and coordinated the work. CEC, RK, and JLR wrote the manuscript and prepared the figures.

## Data and materials availability

Cryo-EM maps are deposited in the Electron Microscopy Data Bank under accession numbers EMD-44350 to 44355. Atomic models are deposited in the Protein Data Bank under accession numbers 9B8O to 9B8Q. Mass spectrometry datasets were deposited to the ProteomeXchange through its partner MassIVE. Assigned accession numbers are MSV000094439, and PXD051100 for the ProteomeXchange.

## List of Supplementary Materials

**1 Supplementary Movie**

**3 Supplementary Figures**

**1 Supplementary Table**

**1 Supplementary Data (Microsoft Excel Spreadsheet)**

**Supplementary Movie 1.** Slices through an electron cryotomogram of SVs on a graphene-oxide coated holey gold grid.

**Supplementary Data 1.** Mass spectrometry of SVs and rat brain lysate.

**Supplementary Figure 1.**
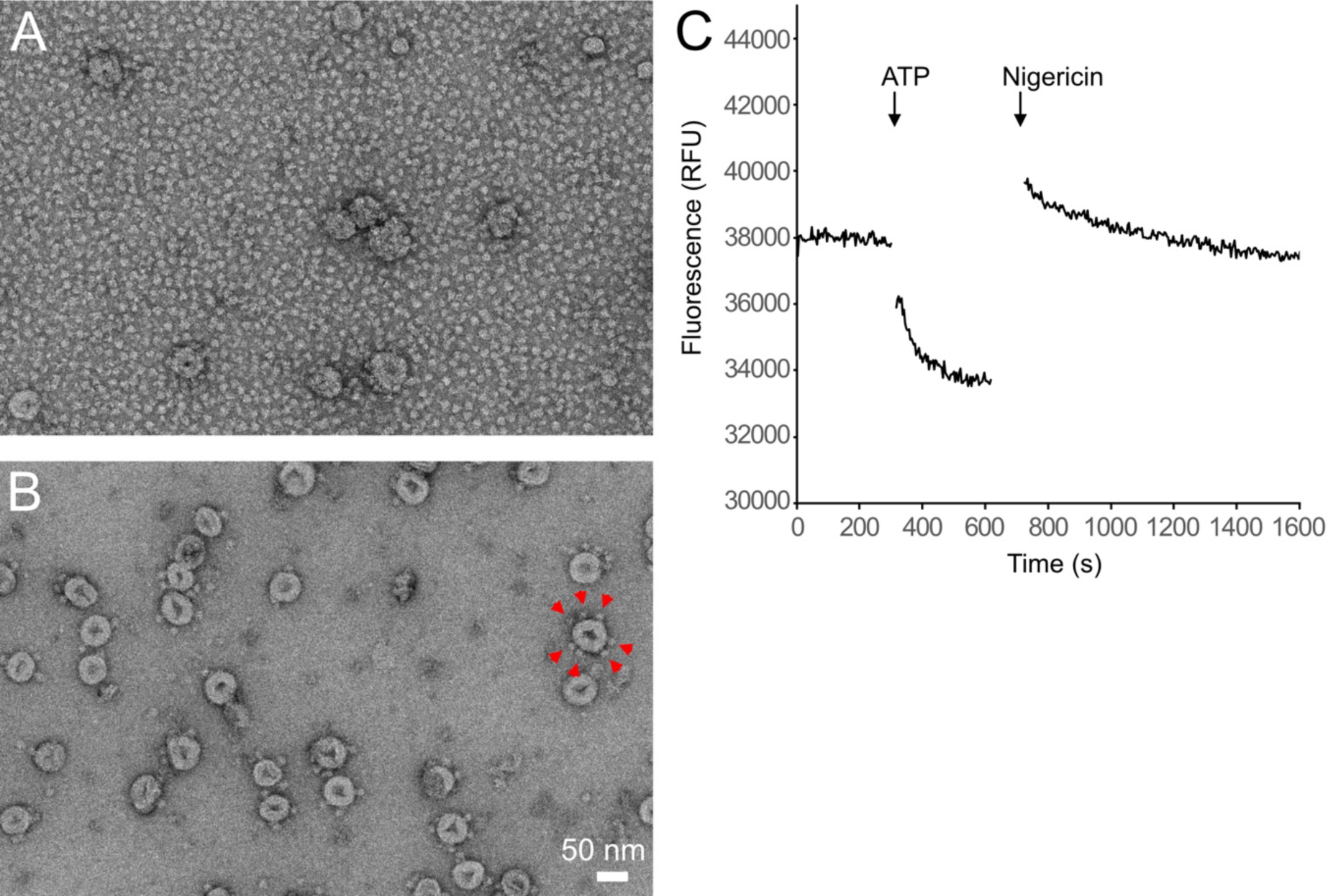
**A,** Negative stain electron micrograph of SVs prior to separation of free V_1_ complexes by gel filtration chromatography. **B,** Negative stain electron micrograph of SVs following separation of free V_1_ complexes by gel filtration chromatography. Red arrows indicate V-ATPase complexes for one of the vesicles. **C,** Acidification of vesicles following gel filtration chromatography. RFU, relative fluorescence units.

**Supplementary Figure 2.**
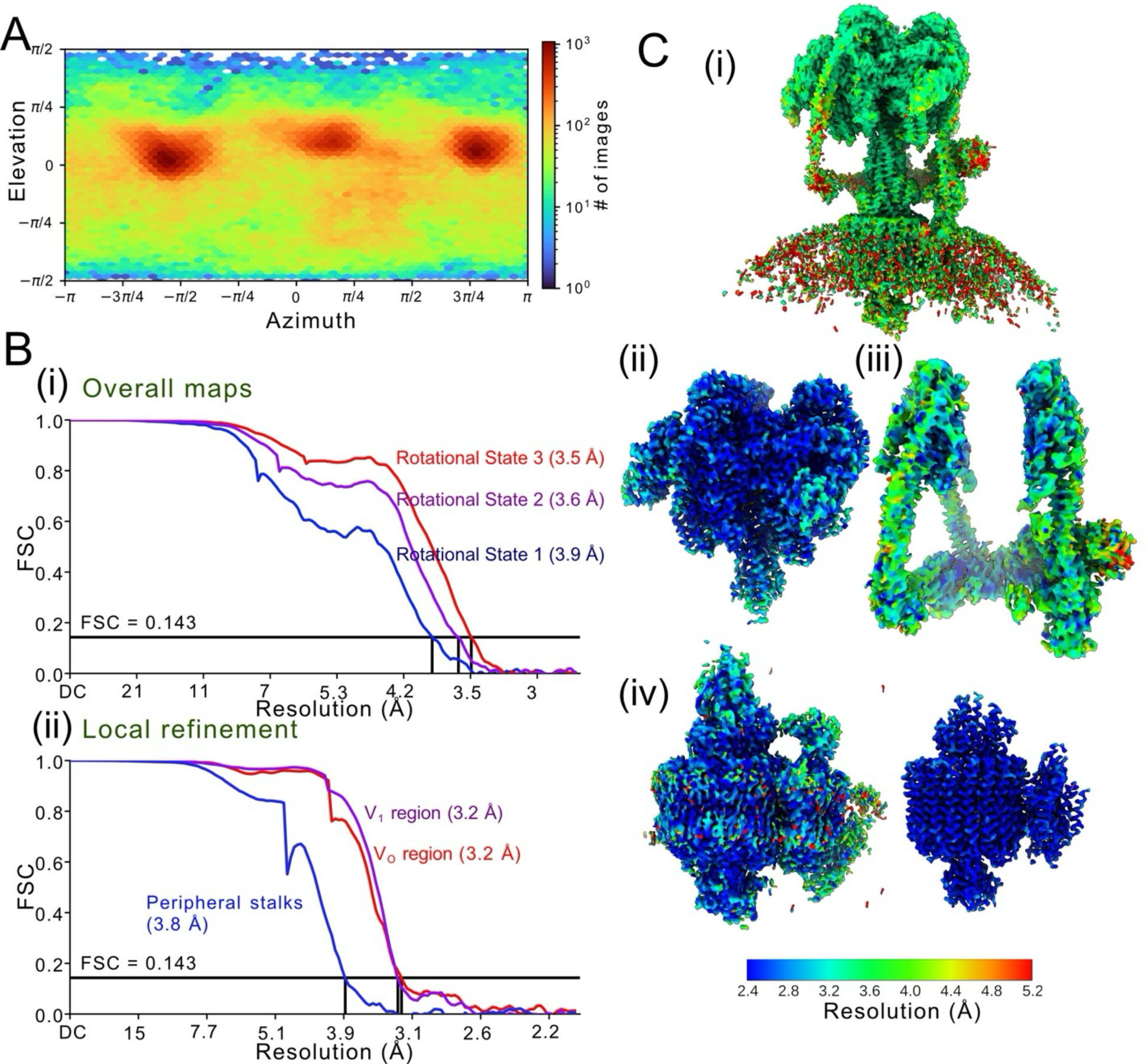
**A,** Orientation distribution plot for particle images of V-ATPase in synaptic vesicles. **B,** Fourier shell correlation (FSC) curves following gold standard refinement with correction for the effects of masking for maps of Rotational States 1, 2, and 3 (**i**) and for local refinement of the V_1_ region, the V_O_ region, and the peripheral stalk structures from Rotational State 3 (**ii**). **C,** Local resolution measurement for the overall map of Rotational State 3 (**i**), the locally refined V_1_ region (**ii**), peripheral stalk structures (**iii**), and V_O_ region at two different density cutoffs (**iv**).

**Supplementary Figure 3.**
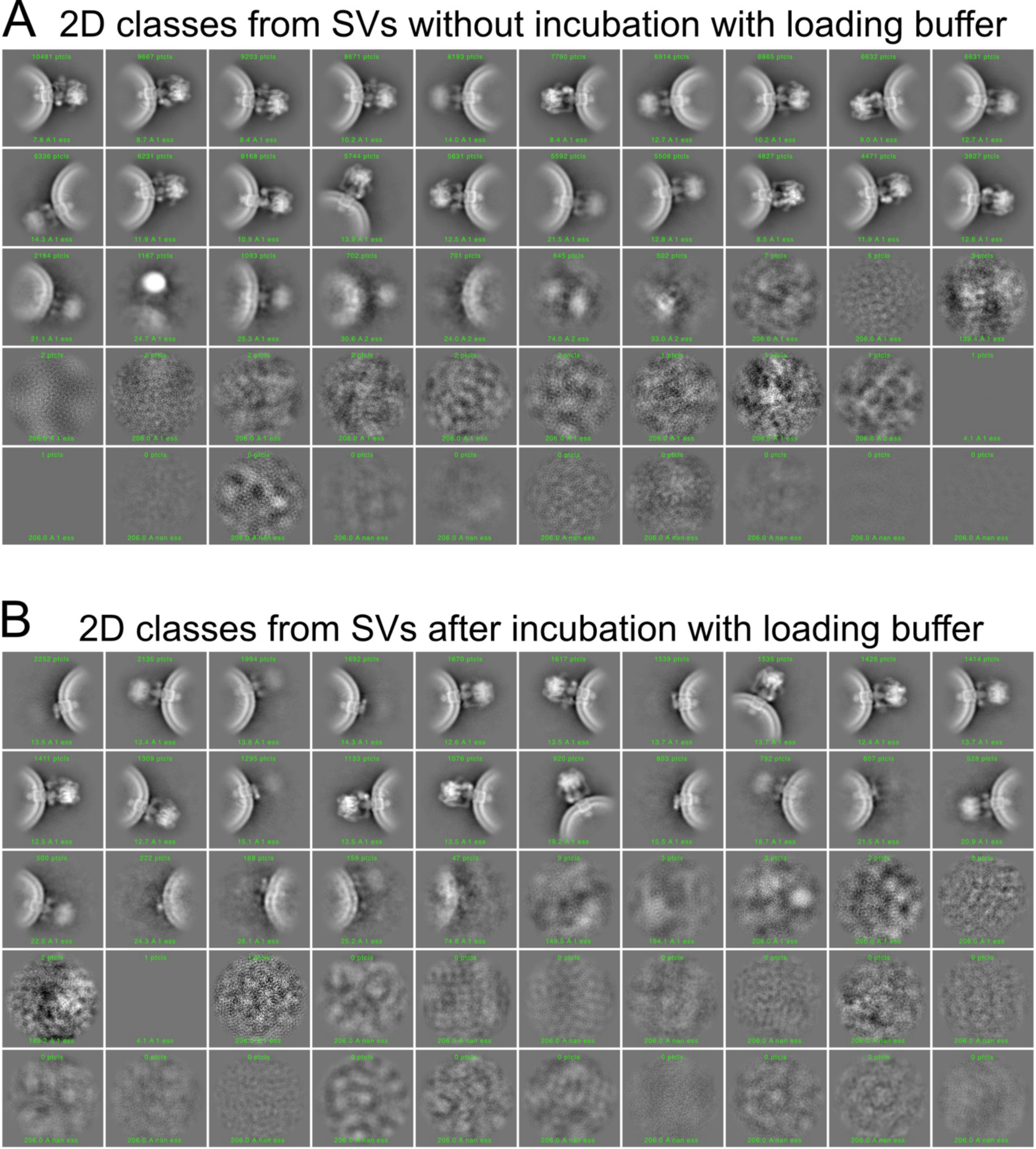
Class averages from 2D classification of exhaustively selected SV edge images before (**A**) and after (**B**) incubation with loading buffer containing 2 mM ATP and 16 mM glutamate.

**Table S1.**
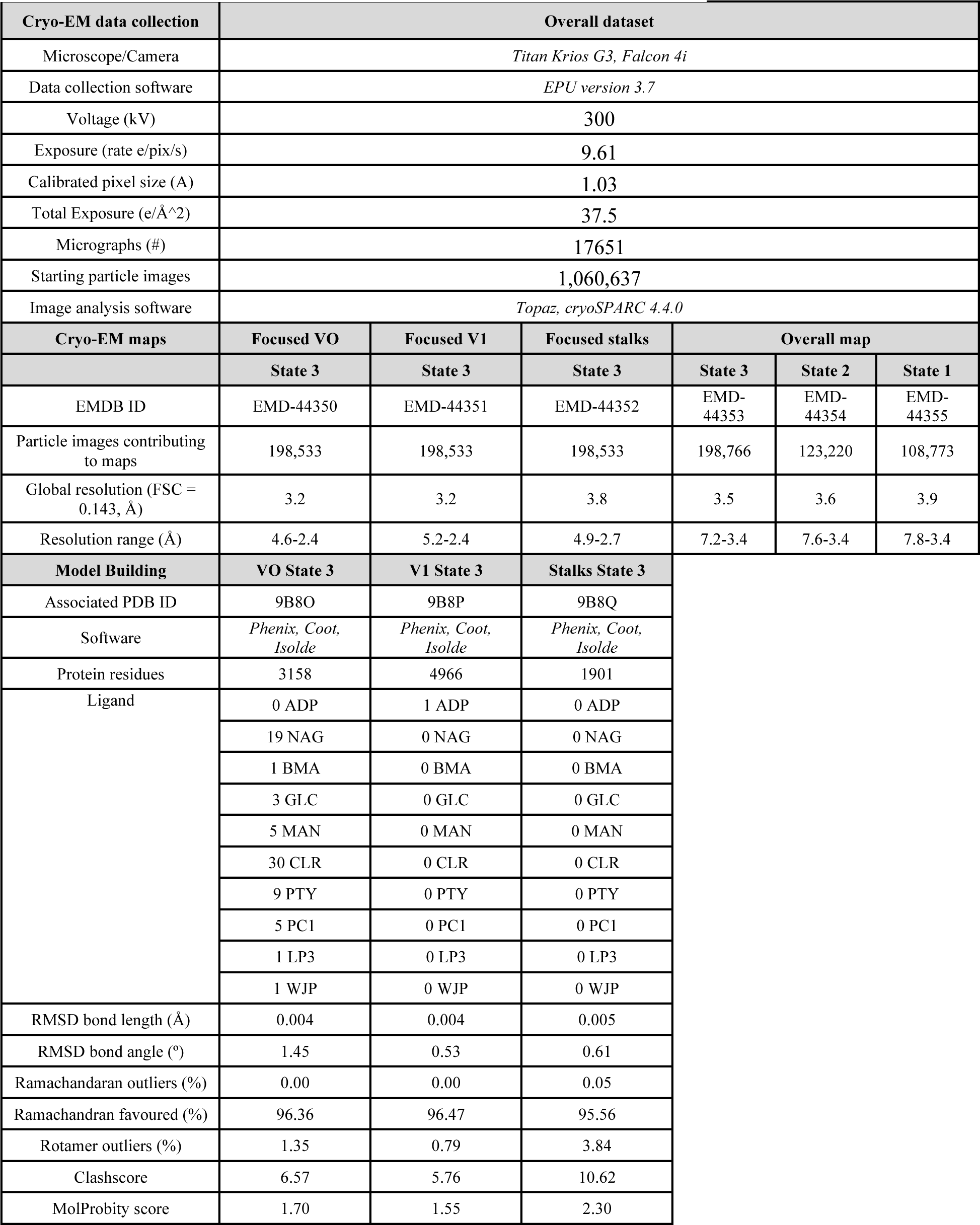
Statistics for cryo-EM and atomic model building.

